# Cytometric fingerprints of gut microbiota predict Crohn’s disease state

**DOI:** 10.1101/649210

**Authors:** Peter Rubbens, Ruben Props, Frederiek-Maarten Kerckhof, Nico Boon, Willem Waegeman

## Abstract

Variations in the gut microbiome have been associated with changes in health state such as Crohn’s disease. Most surveys characterize the microbiome through analysis of the 16S rRNA gene. An alternative technology that can be used is flow cytometry. In this report we analyzed a disease cohort that has been characterized by both technologies. Changes in microbial community structure are reflected in both types of data. We demonstrate that cytometric fingerprints can be used as a diagnostic tool in order to classify samples according to Crohn’s disease state. These results highlight the potential of flow cytometry to perform rapid diagnostics of microbiome-associated diseases.

---

Variations in the gut microbiome have been associated with changes in health state, such as obesity, inflammatory bowel diseases and diabetes (1–3). Characterization of the microbiome is mostly done through analysis of the 16S rRNA gene. Because sequence-based surveys are becoming standardized, microbiome analysis shows great potential to be included in precision medicine (4). Yet, sequence-based surveys are still budget-limited and time intensive (5, 6).

Flow cytometry is a single-cell technology, able to measure up to thousands of individual cells in mere seconds. When applied to microbial communities, both morphological and physiological characteristics are recorded for every cell (7). The aggregation of these cellular characteristics describes the status of a microbial community. By creating a cytometric *fingerprint*, community dynamics and functioning can be quantified and related to space, time or other external variables. Because of the strong connection between the microbiome and human health (8, 9), cytometric fingerprints have the potential to be used as a diagnostic tool to rapidly identify microbiome-associated diseases (10). As such, they have been used to study colitis in murine models (11).

In this study we analyzed the recently published data of a disease cohort containing samples diagnosed with Crohn’s disease (CD) (*n* = 29) and a healthy control (HC) group (*n* = 66). All samples have been analyzed by flow cytometry and 16S rRNA gene amplicon sequencing (12). The original study suggested a clear difference in community structure as measured through 16S rRNA gene sequencing in function of Crohn’s disease. In this work we set forth to demonstrate that these differences are reflected in the cytometry data as well, and in addition, compare the predictive power of both technologies in a straightforward way. We used *PhenoGMM*, which is an adaptive cytometric fingerprinting strategy based on Gaussian Mixture Models, to cluster individual cells in operational groups (13). This results in a contingency table, which stores cell counts per mixture and sample. In other words, cytometric fingerprints result in abundance data. However, abundances are expressed in terms of phenotypically similar cells instead of grouped sequences. Random Forest classification was applied to classify unseen samples according to the disease state based on both types of data (Fig. 1). 16S rRNA gene amplicon sequencing resulted in a perfect identification (mean AUROC = 1.0), with flow cytometry doing marginally worse (mean AUROC = 0.94).

**Fig. 1.**
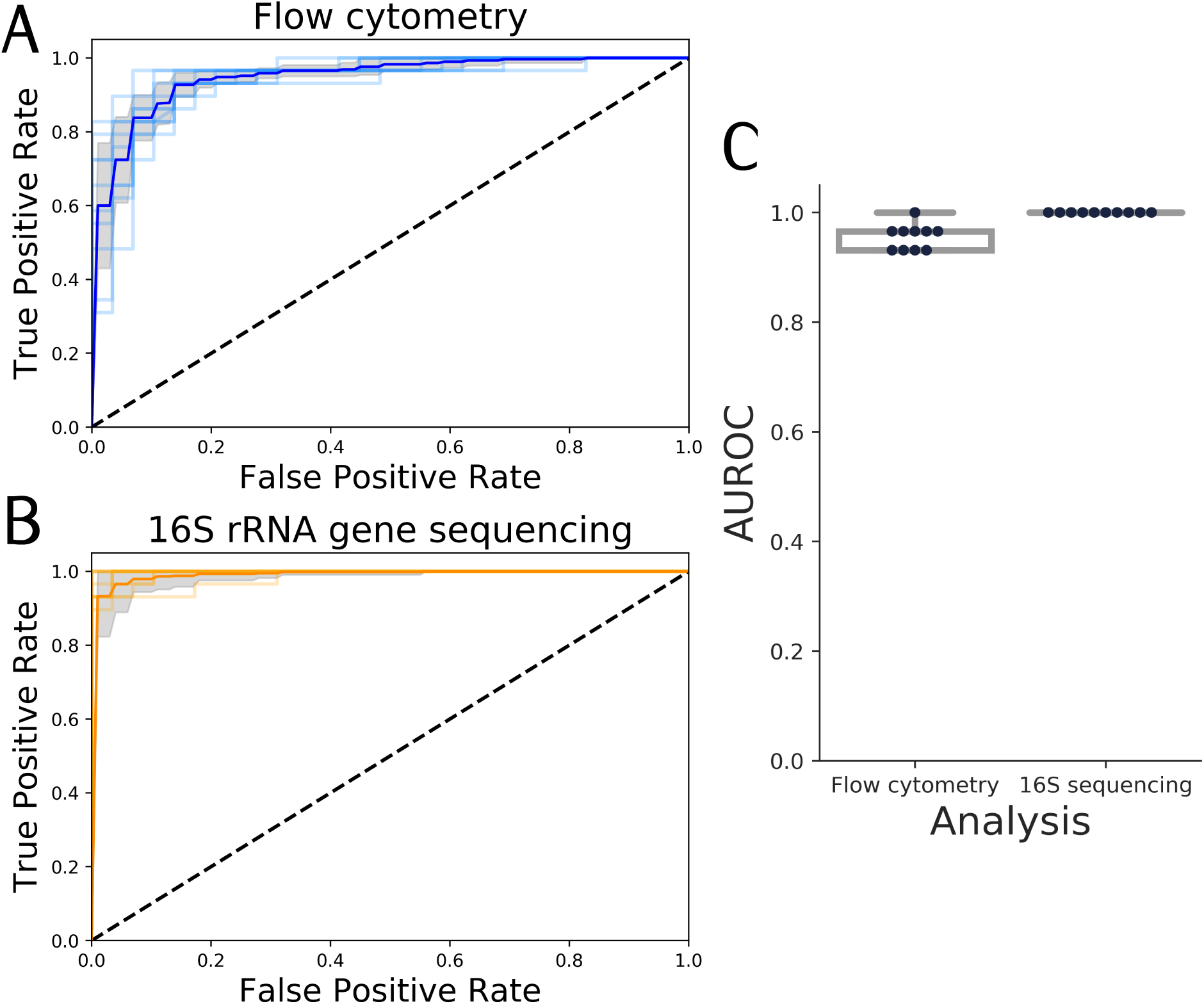
Summary of Random Forest classification of CD vs HC test samples for 10 runs of the model. The test set was created using leave-pair-out cross-validation. **A**: Receiver operating characteristic (ROC) curves, along with the average ROC curve and standard deviation (SD, grey area), based on pooled predictions for cytometry data. **B**: ROC curves, along with the average ROC curve and SD (grey area), based on pooled predictions for genus abundances derived from 16S rRNA gene sequencing data. **C**: Summary of the area under the ROC (AUROC) curves, this time based on pairwise averaging of the test set. Each black dot represents the AUROC for an individual run, along with a visualization of the median. A boxplot displays the first and third quartile and the median line. Whiskers extend from the quartiles to 1.5 times the interquartile range.

The diversity within a microbial community can be used to relate changes in community structure to external gradients or differences between case and control status. Although microbial diversity is mostly assessed through sequencing-based surveys (14), it can also be defined based on flow cytometry data (15). In fact, a number of recent reports have illustrated that the cytometric diversity correlates well with taxonomic diversity based on 16S rRNA gene sequencing (16, 17).

We quantified the within-sample diversity in terms of richness (*D*_0_) and evenness (*D*_2_) for both types of data. Cytometric diversity, based on similarly grouped cells, is significantly correlated with taxonomic diversity based on gene abundances (Spearman’s 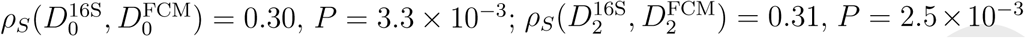). Both taxonomic diversity (Fig 2A) and cytometric diversity (Fig. 2B) were significant markers in function of CD versus HC, in which both the richness and evenness of gut microbiota were significantly lower for CD compared to HC samples (Mann-Whitney-U test: *P <* 1 *×* 10^*-*4^). We also assessed which cytometric groups captured significant changes according to the disease state; 132 contained significantly more cell counts for CD than HC, while 103 groups contained significantly more cell counts for HC than CD (Mann-Whitney U test, *P <* 0.05 after Benjamini-Hochberg correction). The locations of these groups revealed a clear structure (Fig 2C). This proves that structural differences in function of the disease state were captured by the cytometric fingperprints, which can be summarized in terms of the cytometric diversity.

**Fig. 2.**
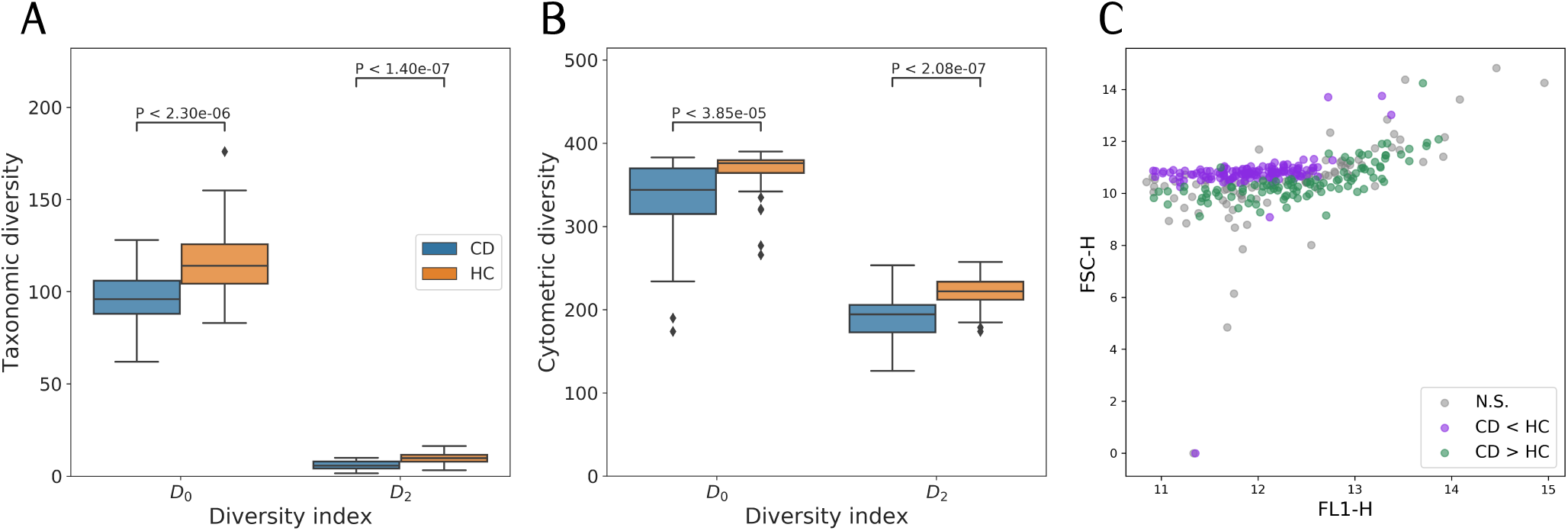
Diversity and cytometric structure for CD (*n* = 29) vs HC (*n* = 66). Statistical differences were assessed using a Mann-Whitney U test. Each boxplot displays the first and third quartile and the median line. Whiskers extend from the quartiles to 1.5 times the interquartile range. Points that lie outside this range are visualized as outliers. **A**, within-sample diversity based on genus abundances derived through 16S rRNA gene amplicon sequencing and **B**, cytometric fingerprinting. **C** Location of the means of each flow cytometric operational group in the FL1-H – FSC-H scatterplot. Groups are annotated whether they contain significantly more cell counts for CD than HC (CD > HC), significantly less cell counts for CD than HC (CD < HC) or whether differences were not significant (NS), *P*-values were corrected by Benjamini-Hochberg correction.

Flow cytometry has become a vital part of clinical diagnostics (18), yet its application to the human microbiome is rarely considered in clinical settings. Although flow cytometry does not allow to inspect the genetic make-up of the microbial community, it may enable rapid and affordable diagnostics of microbiome-associated diseases.

## Materials & Methods

### Dataset description

The data were retrieved from the study by Vandeputte et al. (12). In brief, a disease cohort was analyzed by both flow cytometry and 16S rRNA gene amplicon sequencing. The cohort consists of 29 patients diagnosed with Crohn’s disease (CD) versus 66 healthy control (HC) samples.

### Analysis

Taxa were identified at the genus level based on similar 16S genes, and the genus table was used as reported by Vandeputte et al. Bacterial cells in the fecal stool of each patient were additionally analyzed by flow cytometry. Samples were measured twice at three different days, resulting in six samples per patient. The flow cytometry data were analyzed according to the following steps:

#### Preprocessing

All channels were transformed by *f* (*x*) = asinh(*x*). A fixed gating strategy was used to remove noise in the FL1-SSC space. Additional automated denoising was performed using the FlowAI package (v1.4.4., target channel: FL1, changepoint detection: 150) (19).

#### Cytometric fingerprinting

Cytometric fingerprints were determined using *PhenoGMM* (13). In brief, all samples were first subsampled to the same number of cell counts (288 per sample). Next, samples are split according to a training and test set, after which a Gaussian Mixture Model with 400 mixtures is fitted to the training set, using the FSC-A, SSC-A, FL1-A (488 nm) and FL3-A (610 nm) channels. The fitted Gaussian Mixture Model was used to derive cell counts for the test samples. All available cells were used to store cells per mixture and sample, which resulted in a contingency table of relative cell counts.

#### Patient status classification

The cell and genus abundance tables were used to perform patient status classification (CD versus HC). A test set was created using a leave-pair-out strategy (20), in which one CD sample and one HC were randomly left out. This was repeated 29 times, in such a way that each CD sample was left out once, creating 29 test pairs. The remainder of the samples was used to train a Random Forest classifier with 400 trees (21). The hyperparameters were optimized using a randomized grid search (22). 100 random combinations of hyperparameter values were evaluated using stratified ten-fold cross validation, using the Area under the ROC curve (AUROC) as performance metric. The maximum number of variables that were considered at an individual split for a decision tree was randomly drawn from {1,…, *K*}, in which *K* denotes the number of mixtures or genera, and the minimum number of samples for a specific leaf was randomly drawn between 1,…, 5. Cross-validation, Random Forest classification and performance evaluation were performed using the *scikit-learn* machine learning library (23). ROC curves were created based on pooled predictions of the test set. AUROC values were calculated after averaging predictions per test pair (20).

#### Microbial diversity

The within-sample diversity was calculated based on cell and gene abundances. This was done using the Hill numbers, defined by *D*0 (richness) and *D*2 (evenness) (24). Correspondence between taxonomic diversity and cytometric diversity was assessed using Spearman’s *ρs* correlation with *SciPy*’s *spearmanr()* function (25). Statistical differences between CD and HC cell counts and diversity were assessed according to a Mann-Whitney-U test, using *SciPy*’s *mannwhitneyu()* function. *P*-values were adjusted after Benjamini-Hochberg correction, using the *multipletests()* function from the *statsmodels* package (26).

### Availability

The genus table can be accessed as supporting information to the original publication (12). Denoised flow cytomery data can be accessed via FlowRepository (ID:FR-FCM-ZYVH). Code and data to reproduce the analysis supporting the manuscript can be accessed via https://github.com/prubbens/PhenoGMM_CD.

## ACKNOWLEDGMENTS

We thank Gunther Kathagen and Jeroen Raes for sharing the raw flow cytometry data.

## References

1. PJ Turnbaugh, et al., A core gut microbiome in obese and lean twins. Nature 457, 480–485 (2009).

2. D Gevers, et al., The treatment-naive microbiome in new-onset Crohn’s disease. Cell Host Microbe 15, 382–392 (2014).

3. N Larsen, et al., Gut microbiota in human adults with type 2 diabetes differs from non-diabetic adults. PLoS ONE 5 (2010).

4. TM Kuntz, JA Gilbert, Introducing the Microbiome into Precision Medicine. Trends Pharmacol. Sci. 38, 81–91 (2017).

5. D Sims, I Sudbery, NE Ilott, A Heger, CP Ponting, Sequencing depth and coverage: Key considerations in genomic analyses. Nat. Rev. Genet. 15, 121–132 (2014).

6. J van Dorst, et al., Community fingerprinting in a sequencing world. FEMS Microbiol. Ecol. 89, 316–330 (2014).

7. S Müller, G Nebe-Von-Caron, Functional single-cell analyses: Flow cytometry and cell sorting of microbial populations and communities. FEMS Microbiol. Rev. 34, 554–587 (2010).

8. VB Young, The role of the microbiome in human health and disease: An introduction for clinicians. BMJ 356 (2017).

9. JA Gilbert, et al., Current understanding of the human microbiome. Nat. Medicine 24, 392–400 (2018).

10. C Koch, S Müller, Personalized microbiome dynamics –Cytometric fingerprints for routine diagnostics. Mol. Aspects Medicine 59, 123–134 (2018).

11. J Zimmermann, et al., High-resolution microbiota flow cytometry reveals dynamic colitis-associated changes in fecal bacterial composition. Eur. J. Immunol. 46, 1300–1303 (2016).

12. D Vandeputte, et al., Quantitative microbiome profiling links gut community variation to microbial load. Nature 551, 507–511 (2017).

13. P Rubbens, R Props, FM Kerckhof, N Boon, W Waegeman, PhenoGMM: Gaussian mixture modelling of cytometry data enables efficient predictions of microbial biodiversity. biorXiv, 641464 (2019).

14. SM Gibbons, JA Gilbert, Microbial diversity—exploration of natural ecosystems and microbiomes. Curr. Opin. Genet. & Dev. 35, 66–72 (2015).

15. WKW Li, Cytometric diversity in marine ultraphytoplankton. Limnol. Oceanogr. 42, 874–880 (1997).

16. F. García, L Alonso-Sáez, XA. Morán, Á López-Urrutia, Seasonality in molecular and cytometric diversity of marine bacterioplankton: The re-shuffling of bacterial taxa by vertical mixing. Environ. Microbiol. 17, 4133–4142 (2015).

17. R Props, P Monsieurs, M Mysara, L Clement, N Boon, Measuring the biodiversity of microbial communities by flow cytometry. Methods Ecol. Evol. 7, 1376–1385 (2016).

18. JP Robinson, M Roederer, Flow cytometry strikes gold. Science 350, 739–740 (2015).

19. G Monaco, et al., FlowAI: Automatic and interactive anomaly discerning tools for flow cytometry data. Bioinformatics 32, 2473–2480 (2016).

20. A Airola, T Pahikkala, W Waegeman, B De Baets, T Salakoski, An experimental comparison of cross-validation techniques for estimating the area under the ROC curve. Comput. Stat. Data Analysis 55, 1828–1844 (2011).

21. L Breiman, Random forests. Mach. learning 45, 5–32 (2001).

22. J Bergstra, Y Bengio, Random Search for Hyper-Parameter Optimization. J. Mach. Learn. Res. 13, 281–305 (2012).

23. F Pedregosa, et al., Scikit-learn: Machine Learning in Python. J. Of Mach. Learn. Res. 12, 2825–2830 (2011).

24. MO Hill, Diversity and Evenness : A Unifying Notation and Its Consequences. Ecology 54, 427–432 (1973).

25. E Jones, T Oliphant, P Peterson, et al., SciPy: Open source scientific tools for Python (2001–).

26. S Seabold, J Perktold, Statsmodels: Econometric and statistical modeling with python in 9th Python in Science Conference. (2010).

